# Neuroprotective activity of sub-anesthetic ketamine in a preclinical model of Parkinson’s disease is not blocked by antagonizing brain-derived neurotrophic factor signaling

**DOI:** 10.64898/2026.10.28.762430

**Authors:** Carolyn J. Stopera, Mitchell J. Bartlett, Kelsey Bernard, Rachel Conklin, Sansita Singh, Jennifer B. Frye, Lalitha Madhavan, Kristian R. Doyle, Scott J. Sherman, Kathy Steece-Collier, Torsten Falk

## Abstract

Treatment with low-dose ketamine is neuroprotective in rodent models of acute damage, while high doses of ketamine can be neurotoxic. Our group has previously shown that sub-anesthetic ketamine treatment leads to long-term reduction of L-DOPA-induced dyskinesia (LID) via brain-derived neurotrophic factor (BDNF)-signaling in a rat 6-hydroxydopamine (6-OHDA) model of Parkinson’s disease (PD). In this report, we investigated if sub-anesthetic ketamine is also neuroprotective in a progressive 6-OHDA model of PD via neurotrophic action through the tropomyosin receptor kinase B (TrkB) receptor. We demonstrated that ketamine-treatment (20 mg/kg, i.p.) prior to striatal administration of the neurotoxin 6-OHDA is neuroprotective, providing near complete protection against development of parkinsonian motor deficits. This robust behavioral efficacy occurred in the presence of only modest preservation of tyrosine hydroxylase (TH) striatal levels (∼50-60% loss). Interestingly, based on data from an additional cohort treated with ketamine and ANA-12 (TrkB antagonist) we found that the neuroprotective effects of ketamine are not mediated by TrkB receptors. Semi-quantitative western analysis showed that the relative tyrosine hydroxylase content in the striatum was not significantly increased in ketamine-treated rats compared to vehicle. Unbiased stereology of dopamine neurons in the substantia nigra pars compacta (SNc) revealed no significant differences between any of the groups. The role of BDNF was evaluated with *in situ* hybridization for RNA of *BDNF* and *TrkB*. While *BDNF* expression was not changed, co-administration of ketamine and ANA-12 led to a significant decrease in *TrkB* receptor expression compared to ketamine-only on the lesioned but not the intact hemisphere in the SNc. In conclusion, we show that ketamine at the low dose currently pursued for therapeutic benefit for LID led to robust behavioral motor improvement in this rodent PD degeneration model, yet these protective effects were not driven by neurotrophic BDNF-signaling, indicating complex mechanisms are underlying neuroprotective activities of low-dose ketamine.

## 1. Introduction

Parkinson’s disease (PD) is the second most common neurodegenerative disease, impacting over 1 million people in the United States (Kouli et al., 2018). The cardinal motor symptoms of bradykinesia, resting tremor, rigidity, and postural instability are largely due to the loss of dopaminergic neurons with cell bodies in the substantia nigra (SNc) pars compacta (Kouli et al., 2018). In addition, individuals with PD may also experience akinesia and many other non-motor symptoms such as depression, pain, and cognitive difficulties (Schapira et al., 2017). The current leading pharmacological treatment for PD is the use of dopamine replacement therapy in the form of levodopa (L-DOPA) (Gandhi et al., 2020). However, extended use of L-DOPA eventually leads to the debilitating negative side effects of levodopa-induced dyskinesia (LID) in more than 80% of individuals within 5-10 years of treatment, with this number increasing as the disease progresses (Jankovic, 2005; Cenci, et al. 2010). Therefore, there is a critical need for better management of PD symptoms and development of non-dopaminergic pharmacotherapy treatment options that can extend the therapeutic benefits of L-DOPA and potentially slow disease progression.

Our group has demonstrated that treatment with low-dose, sub-anesthetic ketamine can reduce LID long-term while also providing an acute anti-parkinsonian effect that is independent of L-DOPA administration in a preclinical model (Bartlett et al., 2016; Bartlett et al., 2020; Ye et al., 2021; Stopera et al., 2024; Vishwanath et al., 2025; Falk & Cowen, 2026) and showed further promise of the anti-dyskinetic effects of ketamine in PD patient case reports (Sherman et al., 2016). Sub-anesthetic levels of ketamine have also been effectively used in the treatment of post-traumatic stress disorder (PTSD) and is FDA-approved for treatment-resistant depression (Feder et al., 2014; Walsh et al., 2019; Shiroma et al., 2020). Additionally, sub-anesthetic ketamine treatment has been shown to be neuroprotective in rodent models of cerebral ischemia and traumatic brain injury (Shu et al., 2012; Wang et al., 2017). This neuroprotective activity of low-dose ketamine is in contrast to prior work that showed neurotoxic activity of high anesthetic doses of ketamine indicating dose-specific activities (Chouldhury et al., 2021). Our group specifically has shown that increased brain-derived neurotrophic factor (BDNF) release and activation of its tropomyosin receptor kinase B receptor (TrkB) underlie the long-term anti-dyskinetic activity of ketamine (Bartlett et al., 2020; Bartlett et al., 2025), similar to what was shown for long-term antidepressive activity by ketamine (Casarotto et al., 2021). In other preclinical PD models, an increase in BDNF in the nigrostriatal system has been seen as a result of deep brain stimulation (DBS) in the subthalamic nucleus (STN), as well as the stimulation also having a neuroprotective effect for dopaminergic neurons in the SNc (Maesawa et al., 2004; Spieles-Engemann et al., 2010; Fischer et al., 2017).

Therefore, we hypothesized that sub-anesthetic ketamine treatment may also exhibit neuroprotective effects for dopaminergic neurons in the SNc or at the terminal projections of these neurons in the striatum via neurotrophic effects mediated by the tropomyosin receptor kinase B (TrkB) receptor. In the current study, we produced a mild, progressive model of PD with a mild unilateral intrastriatal lesion of 6-OHDA. We investigated if ketamine at the dose we previously showed to be anti-dyskinetic (20 mg/kg) was neuroprotective using the amphetamine-induced rotation test as a measure of 6-OHDA lesion severity, and if the effects could be blocked via co-administration of ANA-12, an antagonist to BDNF’s receptor TrkB. Additionally, we investigated if ketamine was able to restore motor movement and reduce akinesia associated with PD in the forelimb-adjusting steps (FAS) test and if this effect could also be blocked with ANA-12 administration. Utilizing tissue from the 6-OHDA lesioned animals, we evaluated TH and NeuN co-labeling for an unbiased count of remaining cells in the SNc between each hemisphere. In addition, we performed RNA *in situ* hybridization for *BDNF* and *TrkB* to evaluate target engagement by ANA-12. Finally, we performed semi-quantitative western blot analysis to measure tyrosine hydroxylase (TH) and extracellular signal regulated kinase (ERK) levels in the striatum and SNc.

## 2. Materials & Methods

### 2.1. Abbreviations

In **Table 1** the Abbreviations are introduced.

**Table 1:** Abbreviations.

| Abbreviation | Full name |
| --- | --- |
| 6-OHDA | 6-hydroxydopamine |
| AR | Amphetamine-induced rotation test |
| ARRIVE | Animal Research: Reporting *In Vivo* Experiments guideline |
| BDNF | Brain-derived neurotrophic factor |
| CD68 | Cluster of Differentiation 68 |
| ERK1/2 | Extracellular signal-regulated protein kinases 1 and 2 |
| FAS test | Forelimb adjusting steps test |
| FDA | Food & Drug Administration |
| GFAP | Glial fibrillary acidic protein |
| IBA1 | Ionized calcium binding adaptor molecule 1 |
| *i.p.* | *Intraperitoneal* |
| LID | Levodopa-induced dyskinesia |
| NMDAR | *N*-methyl-D-aspartate receptor |
| PD | Parkinson's disease |
| RPM | Rotations per minute |
| SEM | Standard error of the mean |
| *SNc* | Substantia Nigra pars compacta |
| STR | Striatum |
| TH | Tyrosine Hydroxylase |
| TrkB | Tropomyosin receptor kinase B |

### 2.2. Animals

Adult male, Sprague-Dawley rats (250 g; Inotiv Inc., West Lafayette, IN), were housed two to three per cage throughout the study in a temperature and humidity-controlled room on a 12-hr reverse light/dark cycle. Food and water were provided *ad libitum*. All animal studies were approved by the Institutional Animal Care and Use Committee at The University of Arizona and in accordance with the NIH Guidelines for the Care and Use of Laboratory Animals, and the ARRIVE guidelines.

### 2.3. Unilateral 6-hydroxydopamine (6-OHDA) lesion

Rats (n = 51) were pretreated (*i.p.* 30-min prior to 6-OHDA administration) with 12.5 mg/kg desipramine hydrochloride in 0.9% sterile saline, with 10% dimethyl sulfoxide (DMSO; Sigma-Aldrich), to prevent noradrenergic neuron loss. Isoflurane (VetOne) anesthesia was mixed in a vaporizer (JD Medical, Phoenix, AZ) at 1.5-2.0% with 1.5 L of oxygen per minute. 6-OHDA (13.75 μg in 0.9% sterile saline with 0.02% ascorbic acid; Sigma-Aldrich) was injected (0.5 μl/min) using a microinjector (Stoelting Co., Wood Dale, IL) and 10 μl Hamilton syringe (Model #1701; Hamilton Co., Reno, NV) attached to a 26-gauge needle (Hamilton Co.) into two locations in the striatum: (1) AP +0.2, ML +3.5, DV –6.4 and (2) AP +1.6, ML +2.4, DV –4.0 (Paxinos and Watson, 2007). The syringe remained in place for 5 minutes post-injection to prevent backflow of 6-OHDA solution. Given the established and standard nature of this model both in the field (Galindo et al., 2014; Kaczyńska et al., 2020) and in our laboratory (Bartlett et al., 2016; Bartlett et al., 2020; Ye et al., 2021), no sham-surgery groups were included.

### 2.4. Treatment groups and experimental design

In *Experiment 1*, ketamine (20 mg/kg; VetOne) was formulated in 0.9% USP grade sterile saline (Sigma-Aldrich, St. Louis, MO) and N-[2-[(oxoazepan-3-yl)carbamoyl]phenyl]-1-benzothiopene-2-carboxamide (ANA-12; 0.5 mg/kg; Tocris Bioscience, Minneapolis, MN) was formulated in 5% dimethylsulfoxide (DMSO; Thermo Fisher Scientific, Waltham, MA), 55% polyethylene glycol 400 (PEG-400; Sigma-Aldrich, St. Louis, MO), and 40% sterile saline. All systemic drugs were administered at a volume of 1 ml/kg via *i.p*. injection. Rats were treated 3 times with Vehicle (n = 18), Ketamine (20 mg/kg; n = 16), or Ketamine + ANA-12 (0.5 mg/kg; n = 16) spaced 2 hours apart, beginning 6 hours prior to unilateral 6-OHDA striatal lesion. Rats were subsequently treated with the same 6-hour injection protocol for 7 consecutive days post lesion, as outlined in the Scheme in **Figure 1**. Behavioral assessment was performed at 14-and 28-days post lesion. One animal died due to complications unrelated to the study.

**Figure 1.**
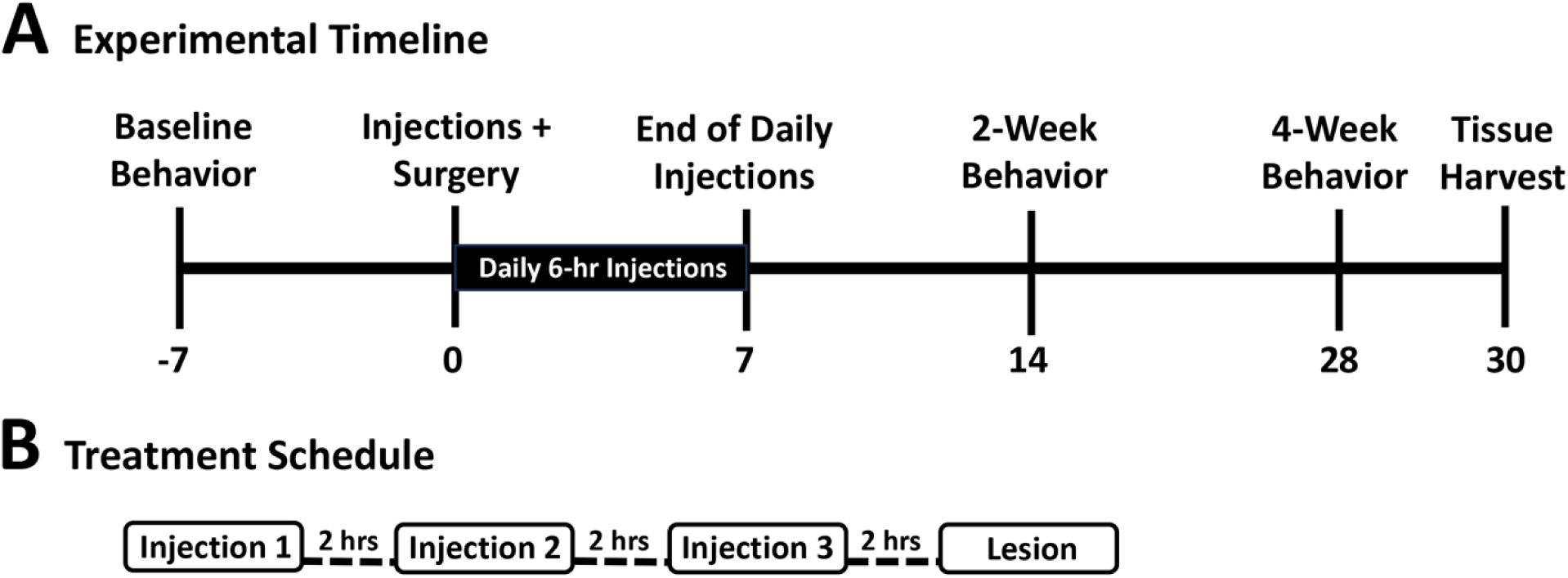
Study scheme for *Experiment*. **(A)** Experimental timeline for the *Experiment*. Injections include daily 6-hour treatment protocol shown in B. **(B)** Treatment injection paradigm utilized before lesion surgery and for 7 consecutive days post-lesion surgery. Injections consisted of vehicle, ketamine (20 mg/kg), or ketamine + ANA-12 (0.5 mg/kg) at each timepoint. ANA-12 was co-administered with ketamine on injection 1 and by itself 1-hour after ketamine injection 3.

### 2.5. Amphetamine-induced rotation test

To verify successful 6-OHDA lesion and determine severity of dopamine neuron denervation after treatment, rats were administered dextro-amphetamine (5 mg/kg in 0.9% sterile saline, *i.p.*; Sigma-Aldrich) and placed into a plexiglass cylinder (9-inch diameter x 12-inch height). Rotational asymmetry was assessed, as done prior (Bartlett et al., 2016; Bartlett et al., 2020), by counting net rotations ipsilateral to the lesion every minute over a 90-minute period at 14– and 28-days post-lesion.

### 2.6. Forelimb adjusting steps (FAS) test

The forelimb-adjusting steps (FAS) test is used to measure akinesia, a common symptom of PD (Bartlett et al., 2020, Lindenbach et al., 2011). In each trial, both hindlimbs and one forepaw were restrained by the investigator and the rat was moved laterally in either direction with the free forepaw supporting its weight. Trials consisted of moving the rat across a 90 cm distance at a rate of 9 cm/second. All trials were video recorded and the mean ± SEM number of steps were counted.

### 2.7. Tissue preparation for western blot

Rats were euthanized 30 days post-lesion with carbon dioxide. Whole striatum and SNc tissue was dissected from both brain hemispheres and flash frozen in liquid nitrogen. Tissue was homogenized in cold phosphate-buffered saline containing 1% Triton X-100, 0.1% sodium dodecyl sulfate, protease inhibitor cocktail tablet (ThermoFisher Scientific, Waltham, MA) and phosphatase inhibitor cocktail tablet (ThermoFisher Scientific). The samples were centrifuged (Model, Location) at 4°C for 30 minutes at 14,000 x g. Protein concentration for each sample was determined using a Pierce^TM^ Rapid Gold BCA Protein Assay Kit (ThermoFisher Scientific) and read on a Flexstation 3 (Molecular Devices, San Diego, CA).

### 2.8. Western blot analysis

All tissue samples were run on Any-kD MiniProtean^®^TGX™ (Bio-Rad Laboratories, Hercules, CA) precast gels and transferred with a Trans-Blot TurboTransfer System (Bio-Rad Laboratories). A Chameleon® Duo Pre-stained Protein Ladder (LI-COR, Lincoln, NE) was used for molecular weight reference. Blots were scanned on an Odyssey CLx imaging system (LI-COR) and analyzed with Empiria Studio Software (LI-COR). Striatal samples were loaded into each gel at a concentration of 1 μg/μL with 4.1 μg of total protein. SNc samples were loaded into each gel at a concentration of 3 μg/μL with 17 μg of total protein. Primary antibodies: Tyrosine Hydroxylase (TH, 1:2000; Millipore, Billerica, MA), Beta-Actin (1:10000; Sigma-Aldrich), Phospho-p-44/42 MAPK (Erk1/2) (1:2000; Cell Signaling Technologies; Danvers, MA), p-44/42 MAPK (Erk1/2) (1:2000; Cell Signaling Technologies; Danvers, MA). Secondary antibodies: IRDye® 800 CW Donkey anti-rabbit (1:20000; LI-COR), IRDye® 680 CW Donkey anti-mouse (1:20000; LI-COR).

### 2.9. Immunohistochemistry for neuronal stereology

Fixed tissue was sectioned using a cryostat (CM1850, Leica, Wetzlar, Germany) at 40 μm. Coronal tissue slices were used for immunohistochemistry with primary antibodies: TH (1:4000; Millipore, Billerica, MA) and NeuN (1:6000; Millipore, Billerica, MA). Secondary antibodies: (Vector Laboratories, Newark, CA). Tissue was stained using Vector Blue Substrate Alkaline Phosphatase Kit and Vector NovaRED Substrate Peroxidase Kit (Vector Laboratories; Newark, CA).

### 2.10. Unbiased neuronal stereology

Unbiased stereology was performed by a blinded investigator. TH and NeuN positive cells were counted in the SNc of each hemisphere in 8 coronal tissue slices per animal. Stereological probes were placed using StereoInvestigator software (v2019.1.3; MBF Bioscience, VT, USA) on a Zeiss Imager M2 microscope (Carl Zeiss, Jena, Germany). Cells were counted using a 170 x 100 μm grid and 75 x 75 μm counting frame. SNc regions were outlined with a 2.5x objective and cells were counted with a 63x oil immersion objective.

### 2.11. In-situ hybridization

Sections were removed from cryoprotectant and rinsed with 0.05M tris-buffered saline (TBS), incubated in ACD hydrogen peroxide solution (Advanced Cell Diagnostics, Newark, CA) for 15 minutes, rinsed again with TBS then mounted on Fisherbrand™ Superfrost™ Plus Microscope Slides and dried on a slide warmer at 60℃ for 24 hours. Slides were then incubated in 100℃ Target Retrieval Solution (Advanced Cell Diagnostics, Newark, CA) for 30 minutes followed by 3 dips in dH2O then 1 minute in 100% ethanol. After slides air-dried at room temperature (RT), tissue sections were outlined with ImmEdge PAP pen (Vector Labs, Burlingame, CA) and left to dry for 10 min at RT before covering each section with Protease III (Advanced Cell Diagnostics, Newark, CA) and placed in 40℃ hybridization oven for 30 minutes. Probe 1 was diluted in probe 2 and the subsequent solution was dropped on tissue and incubated in hybridization oven for 2 hours at 40℃. Amplification steps from the RNAscope Multiplex Fluorescent v2 kit (Advanced Cell Diagnostics, Newark, CA) were followed according to manufacturer’s instructions and developed with Opal™ 690 for TrkB RNA 620 for BDNF RNA diluted in ACD proprietary diluent (Advanced Cell Diagnostics, Newark, CA). After development tissue was further stained for TH (MAB318; Millipore Sigma, Burlington, MA) by first incubating slides in blocking buffer (10% Normal Goat Serum+TBS) for 45 minutes at room temperature then in primary solution (5% Normal Goat Serum+TBS+Mouse anti-TH, MAB318; Millipore Sigma, Burlington, MA) overnight at room temperature. Slides were then rinsed with TBS 3 times followed by incubation in secondary solution (5% NGS+TBS+Alexafluor 488; Advanced Cell Diagnostics, Newark, CA) for 2 hours at RT. Slides were rinsed again with TBS and coverslipped with Vector Hardset with Dapi (Vector H-1500; Vector Laboratories, Newark, CA).

Z-stacks (2um steps) were taken on a Nikon Eclipse Ni microscope (Nikon Instruments; Melville, NY) and captured using NIS Elements software (Nikon Instruments; Melville, NY). Three stacks were acquired to capture the entire SNc (both hemispheres) using the 20x objective and keeping identical settings across all images. Confocal z-stacks were imported into the image visualization and analysis software, Imaris® (v. 9.8.0, Oxford Instruments, Abington, United Kingdom) The surface function was used with semi-automatic thresholding to create 3D reconstructions of TH+ cells. The spots function was used to visualize BDNF mRNA and TrkB mRNA puncta in each image. Parameters for spots were the same for all images. The MATLAB® “Split into Surface Objects” plugin was then applied to the *BDNF and TrkB* mRNA spots. With the plugin we were able to determine the number of BDNF and TrkB “spots” within the TH+ cells, along with the total number of TH+ cells, TrkB and BDNF spots.

### 2.12. Statistical analyses

Statistical analyses were performed using GraphPad Prism version 9.5.1 software (GraphPad Software, La Jolla, CA). The null hypothesis was rejected when p < 0.05. All values are represented as mean ± SEM. Amphetamine-induced rotation tests were analyzed with one-way ANOVAs and Fisher’s LSD *post hoc* tests. FAS tests were analyzed with one-way ANOVAs and Tukey-Kramer corrected *post hoc* tests. Western blot analyses were performed with one-way ANOVAs and Fisher’s LSD *post hoc* tests on data before normalization to vehicle control. The western blot analyzing pERK1/2 was normalized to total ERK1/2. Stereology was analyzed One-way ANOVA and Tukey multiple comparisons *post hoc* tests. RNA *in-situ* hybridization was analyzed with *t*-tests and ANOVAs.

## 3. Results

### 3.1. Sub-anesthetic ketamine treatment provides protection against parkinsonian motor deficits in the progressive 6-OHDA model of PD and this is not blocked by TrkB antagonist ANA-12

Unilateral intrastriatal 6-OHDA lesions generate a mild, progressive model of PD (Mendes-Pinheiro et al., 2021). Lesion severity after 6-OHDA injection can be verified via the amphetamine-induced rotation test as a measure of rotational asymmetry counting the number of net ipsilateral rotations toward the lesion made by the animal. Rats were injected with D-amphetamine (5 mg/kg *i.p.*) at 14– and 28-days post-lesion and rotational asymmetry was assessed. At 14-days post-lesion **(Figure 2A)**, ketamine (K; 169.50 ± 84.35) treated animals showed a decreased number of net ipsilateral rotations compared to vehicle (V; 434.4 ± 155.80) treated animals (One-way ANOVA, Fisher’s LSD *post hoc* tests, p = 0.0827; n = 16-18/group). Ketamine + ANA-12 (K+ANA-12; –21.03 ± 23.95) treated animals showed a decreased number of net ipsilateral rotations compared to ketamine treated animals (One-way ANOVA, Fisher’s LSD *post hoc* tests, p = 0.2149; n = 16/group). Ketamine + ANA-12 treated animals also showed a significantly decreased number of net ipsilateral rotations compared to vehicle treated animals (One-way ANOVA, Fisher’s LSD *post hoc* tests, **p = 0.0038; n = 16-18/group). At 28-days post-lesion **(Figure 2B)**, ketamine (K; 199.5 ± 109.0) treated animals showed a significantly decreased number of net ipsilateral rotations compared to vehicle (V; 582.2 ± 173.4) treated animals (One-way ANOVA, Fisher’s LSD *post hoc* tests, p = 0.0309; n = 16-18/group). Ketamine + ANA-12 (K+ANA-12; –17.09 ± 26.43) treated animals showed a trend of an even further decreased number of net ipsilateral rotations compared to ketamine treated animals (One-way ANOVA, Fisher’s LSD *post hoc* tests, p = 0.2208; n = 16/group). Ketamine + ANA-12 treated animals also showed a significantly decreased number of net ipsilateral rotations compared to vehicle treated animals (One-way ANOVA, Fisher’s LSD *post hoc* tests, **p = 0.0011; n = 16/group).

**Figure 2.**
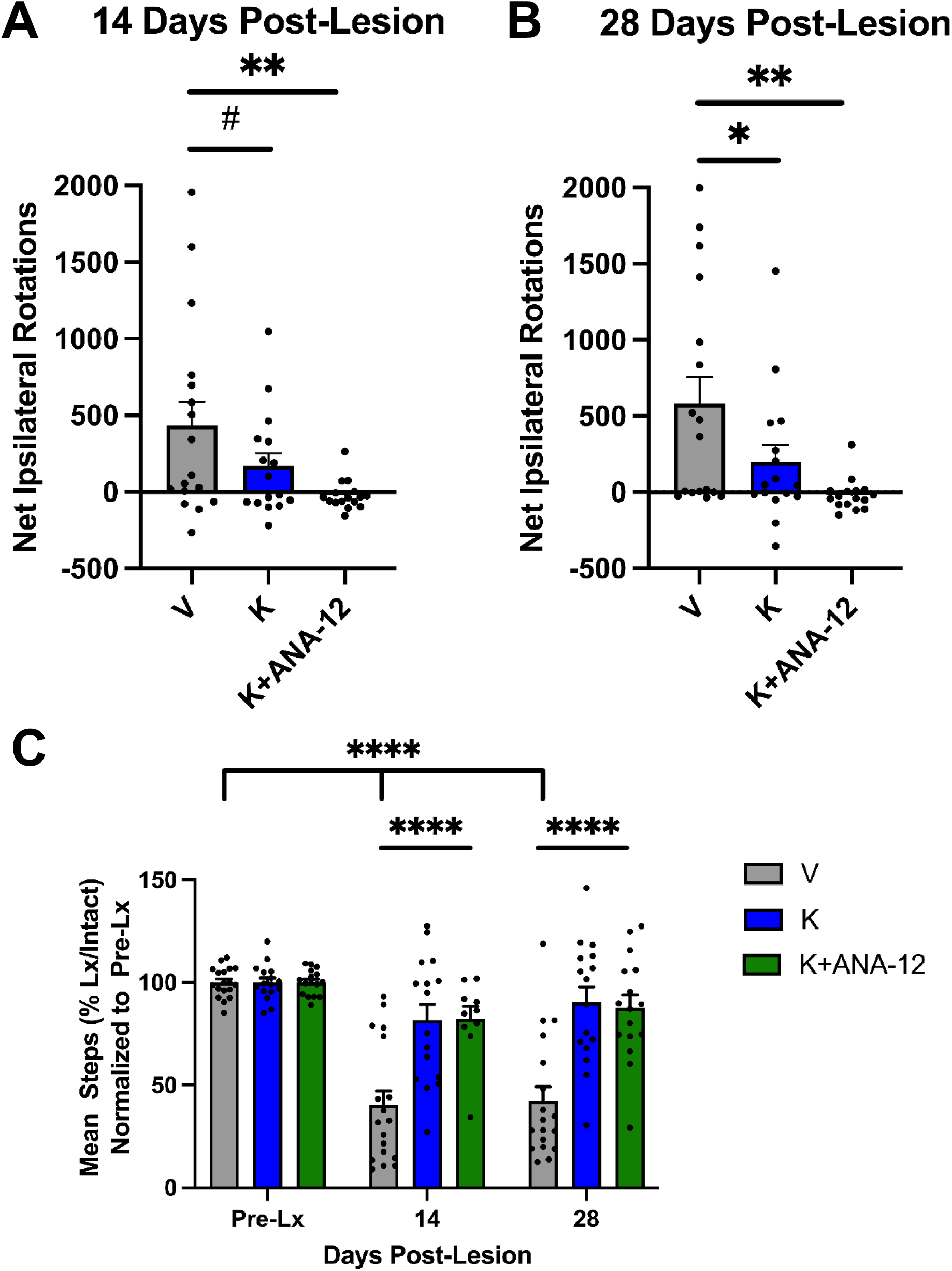
Sub-anesthetic ketamine treatment provides near complete protection against parkinsonian motor deficits in the progressive 6-OHDA model of PD. Rotational asymmetry was assessed at 14– and 28-days post unilateral 6-OHDA striatal lesion by counting the net ipsilateral rotations (mean ± SEM) over 90-minutes. (**A**) Ketamine (K) reduced net ipsilateral rotations at 14-days post-lesion compared to the vehicle group. Ketamine + ANA-12 treated animals showed an even fewer number of net ipsilateral rotations compared to both ketamine and vehicle treated animals. (**B**) Ketamine (K) treated rats showed a decreased number of net ipsilateral rotations at 28-days post-lesion compared to the vehicle (V); group. Ketamine+ANA-12 showed an even fewer number of net ipsilateral rotations compared to ketamine treated animals and a significantly decreased number of net ipsilateral rotations compared to vehicle treated animals. One-way ANOVAs, Fisher’s LSD post hoc tests; *p < 0.05, **p < 0.01, **^#^**p = 0.08; n = 16-18/group. (**C**) Sub-anesthetic ketamine treatment increased number of forelimb adjusting steps (FAST) in the 6-OHDA model of PD. At the 14 and 28-days post-lesion timepoint, ketamine-treated rats (n = 16) showed a significantly increased number of steps made by the weight-bearing forelimb compared to the vehicle group. Ketamine+ANA-12 treated animals showed similar mean steps as the ketamine treated group and showed a significantly increased number of steps compared to the vehicle group. One-way ANOVAs; ****p < 0.0001; n = 16-18/group.

The forelimb adjusting steps (FAS) test is used as a measure of parkinsonian akinesia. The FAS test was performed at 14– and 28-days post-lesion. At 14-days post lesion **(Figure 2C)**, ketamine treated rats showed a significantly increased number of steps (81.70 ± 7.60) compared to the vehicle group (40.30 2± 1.72) (****p < 0.0001, one-way ANOVA, Tukey multiple comparisons *post hoc* tests). At 28-days post-lesion **(Figure 2C)**, ketamine treated rats (90.37 ± 7.45) showed an even greater increase in number of steps compared to the vehicle group (42.35 ± 6.94) (****p < 0.0001, one-way ANOVA, Tukey multiple comparisons *post hoc* tests) indicating a protective activity by ketamine using this non-drug-induced test. The rats treated with Ketamine+ANA-12 showed a similar mean number of steps as the ketamine group at 14-days post-lesion (82.35 ± 6.05) and 28-days post-lesion (87.54 ± 6.34) and showed a significantly increased number of steps compared to the vehicle treated animals (****p < 0.0001, one-way ANOVA, Tukey multiple comparisons *post hoc* tests), further indicating that the TrkB receptor is not involved in the protective activity of ketamine.

### 3.2 Ketamine does not significantly protect against dopaminergic neuron death in the substantia nigra

Substantia nigra tissue from each group was co-labeled with tyrosine hydroxylase (TH) to stain dopaminergic neurons and neuronal nuclei (NeuN) to stain all residing neurons. The number of dopaminergic neurons remaining in the intact and lesioned hemispheres were counted. There were no significant differences between number of dopaminergic neurons in the intact hemisphere between any treatment group (Vehicle: 12401 ± 803.78; Ketamine: 13636.87 ± 804.01; Ketamine + ANA-12: 11580.37 ± 485.77, n = 8/group; **Figure 3A-D**). There was a significant decrease in number of dopaminergic neurons in the lesioned hemisphere compared to the intact hemisphere for all groups (Vehicle: ****p < 0.0001; Ketamine: ****p < 0.0001; Ketamine + ANA-12 **p = 0.0028, n = 8/group). While the number of remaining dopaminergic neurons after ketamine was unchanged, there was a trend of an increased survival (∼30%) in the ketamine + ANA-12 group when compared to the vehicle group, those increases were not significant in the lesioned hemisphere (Vehicle: 4698.75 ± 765.42; Ketamine: 5393 ± 910.17; Ketamine + ANA-12: 6757 ± 1144.49, n = 8/group). One-way ANOVAs with Tukey multiple comparisons post hoc tests.

**Figure 3.**
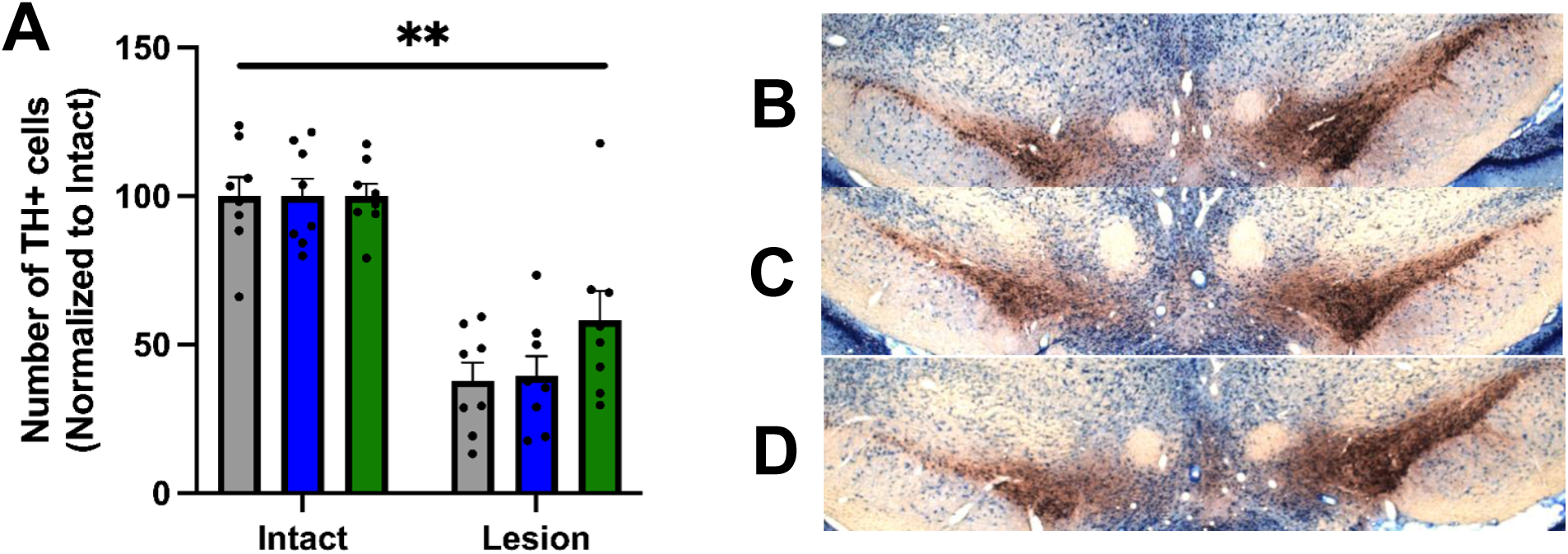
Stereological dopaminergic neuron count in the SNc after parkinsonian lesion. Example photomicrographs of tyrosine hydroxylase (TH; red) and NeuN (blue) co-labeling in the *substantia nigra* of 6-OHDA lesioned animals. (**A**) While there was a trend of ∼30% increased survival of dopaminergic neurons in the ketamine + ANA-12 group the differences were not significant (n=8). One-way ANOVAs, Tukey multiple comparisons *post hoc* tests, **p<0.01. (**B**) TH and NeuN co-labeling in a vehicle treated animal. (**C**) TH and NeuN co-labeling in a ketamine treated animal. (**D**) TH and NeuN co-labeling in a ketamine + ANA-12 treated animal.

### 3.3 Ketamine treatment increases total neurotrophic receptor tyrosine kinase 2 (Ntrk2) expression but not total brain derived neurotrophic factor (BDNF)

*Substantia nigra* tissue was probed for *BDNF* RNA and *TrkB* RNA in the ipsilateral and contralateral hemispheres **(Figure 4)**. *In-situ* hybridization revealed that ketamine + ANA-12 co-treatment significantly decreased total *Ntrk2*, the gene that encodes TrkB, on both the ipsilateral hemisphere (*p = 0.0135, n = 8/group) and the contralateral hemisphere (*p = 0.0254, n = 8/group) compared to the ipsilateral hemisphere of the ketamine treated group (one-way ANOVA, Tukey multiple comparisons *post hoc* tests; **Figure 4A**). This indicates target engagement of a blockade of the TrkB receptor with receptor antagonist ANA-12. There were no significant differences in total *BDNF* between any of the treatment groups or hemispheres **(Figure 4B)**. One-way ANOVAs, Tukey multiple comparisons *post hoc* tests. Example photomicrographs for the 4 conditions are displayed in the panels in **Figure 4C-F**.

**Figure 4.**
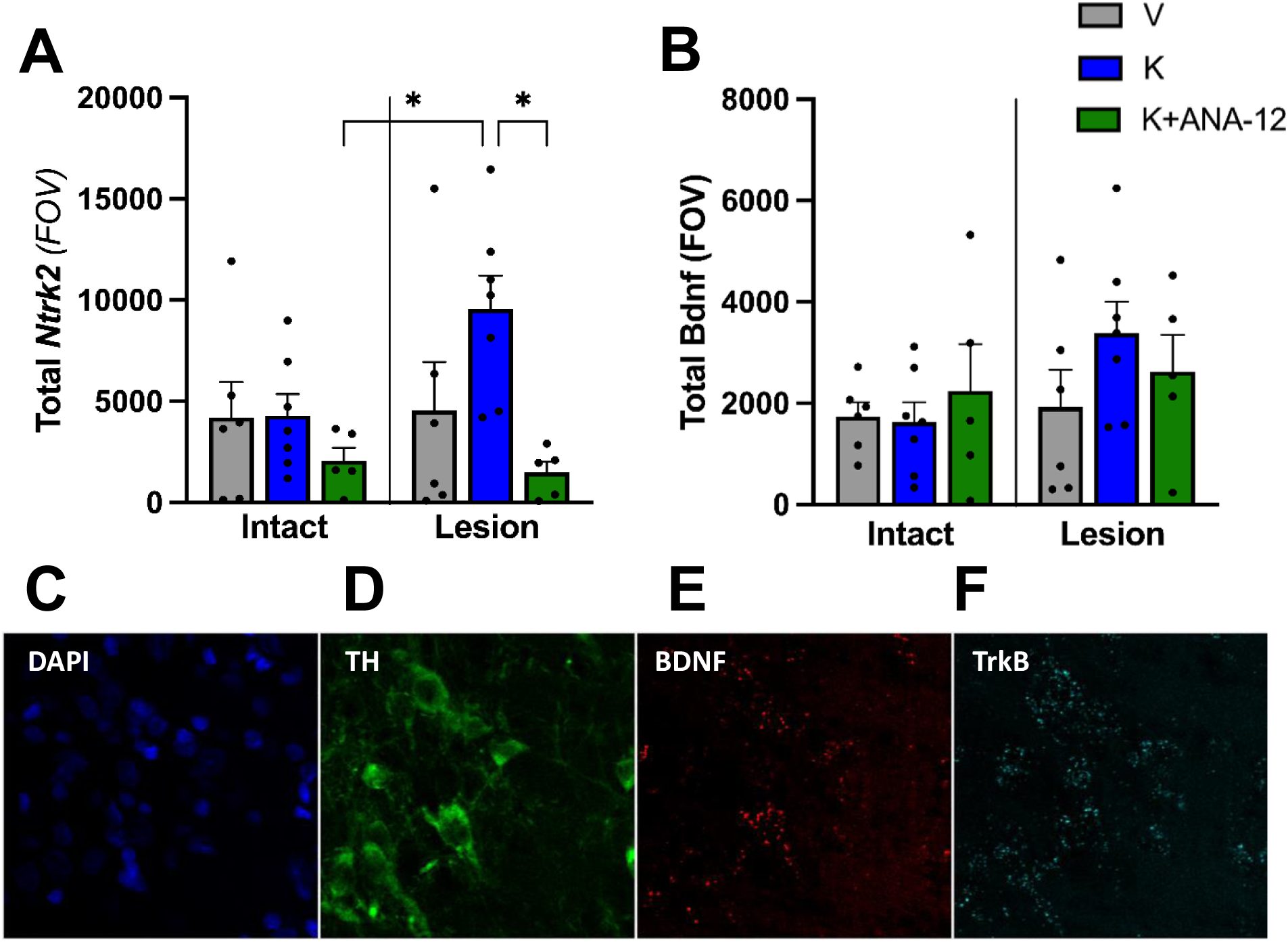
ANA-12 blocks ketamine-induced increase in nigral *TrkB*. (**A**) The mean (± SEM) total *Ntrk2* RNA in the intact and lesioned hemispheres of the SNc is depicted. The data shows that ANA-12 treatment completely blocked a ketamine induced increase of TrkB on the lesioned side. (**B**) The mean total *BDNF* RNA is unchanged in the intact and lesioned hemispheres of the SNc. One-way ANOVA, Tukey multiple comparisons *post hoc* tests; *p<0.05; n=8/group. Example photomicrographs of RNA *in-situ* hybridization for *BDNF* and *TrkB* are provided. Example images are shown for (**C**) DAPI, (**D**) TH, (**E**) *BDNF* RNA, (**F**) *TrkB* RNA.

### 3.4 Evaluation of severity of striatal unilateral 6-OHDA lesion and of phospho-ERK1/2 levels

Semi-quantitative western analysis was performed on striatal tissue measuring TH. The mean ± SEM striatal TH expression in the intact and lesioned hemispheres were evaluated by examining the ratio of percent of TH compared to the internal standard B-actin, with data normalized to the intact vehicle group. While the small increase in TH level vs. vehicle control on the lesioned side was not significance, there is a significant increase in striatal TH levels (mean ± SEM) in the ketamine + ANA-12 co-treated group compared to both the vehicle (27%) and ketamine (18%) treated groups (n=6-8/group, **Figure 5A**). Analysis was performed on raw data prior to normalization.

**Figure 5.**
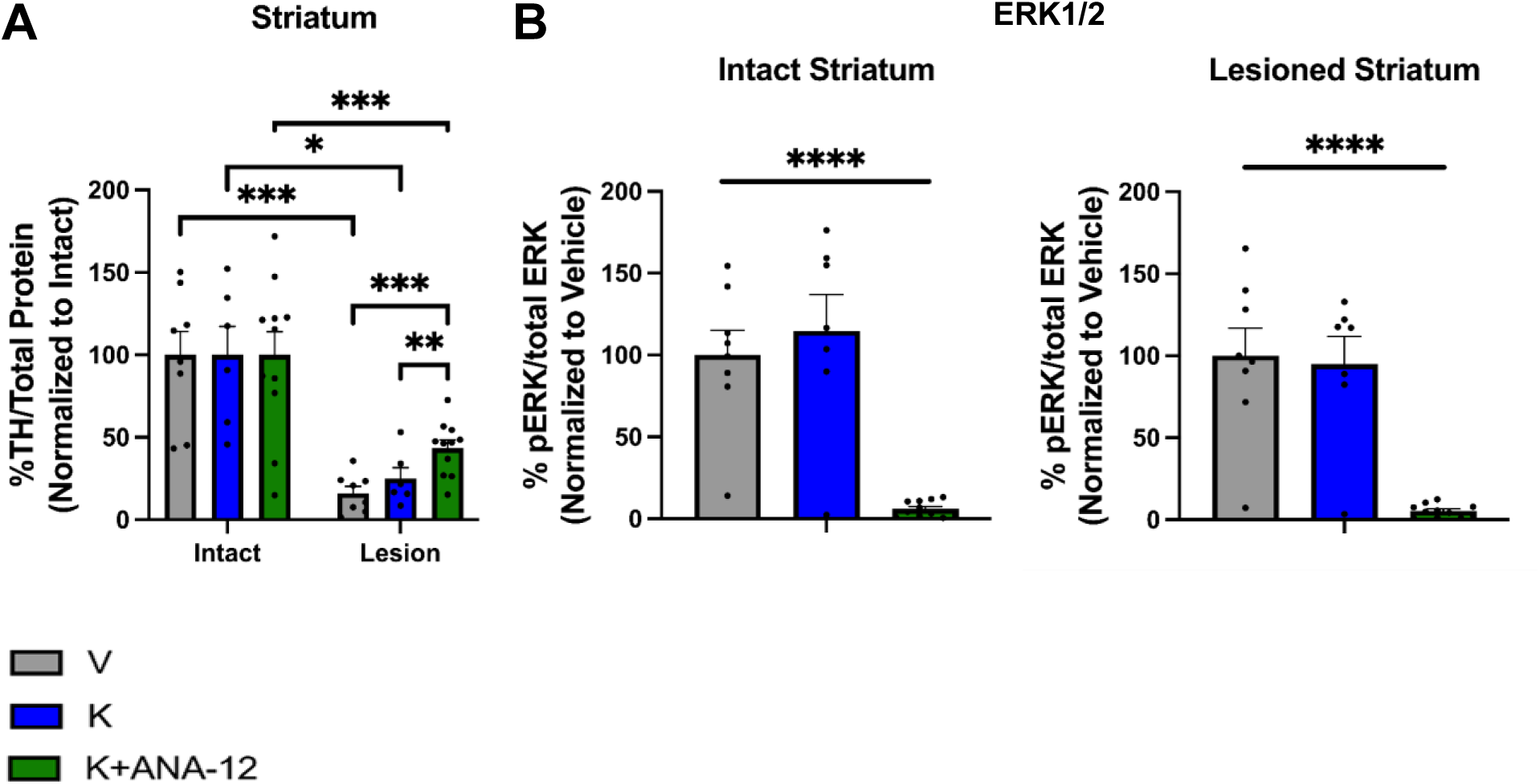
Semi-quantitative measurement of TH and ERK1/2 in striatum. (**A**) There is a trend of increased striatal TH level (mean ± SEM) post-ketamine vs. vehicle, and a significant increase in striatal TH level in the ketamine + ANA-12 co-treated group compared to both the vehicle and ketamine treated groups (n=6-8/group). One-way ANOVA, Tukey multiple comparisons post hoc tests. Analysis performed on data before normalization, *p<0.05, **p<0.001, ***p<0.001. (**B**) There is a significant decrease in p-ERK1/2 levels in the intact and in the lesioned striatum, indicating target engagement of ANA-12, as ERK1/2 is a downstream target of TrkB. One-way ANOVA, Tukey multiple comparisons post hoc tests. ****p<0.0001, n=8/group. Analysis was performed on data before normalization.

Semi-quantitative western analysis was also performed on striatal tissue measuring p-ERK1/2. The mean ± SEM striatal p-ERK1/2 expression in the intact and lesioned hemispheres were evaluated by examining the ratio of percent of p-ERK1/2 compared to total ERK1/2, with data normalized to the vehicle group. There was a significant decrease in p-ERK1/2 expression in the Ketamine + ANA-12 treated group compared to both vehicle and ketamine on the intact hemisphere (****p < 0.0001, n = 8/group; **Figure 5B**) and lesioned hemisphere (****p < 0.0001, n = 8/group; **Figure 5B**). One-way ANOVA, Tukey multiple comparisons post hoc tests. Analysis was performed on raw data prior to normalization.

## 4. Discussion

Ketamine is widely known to be an effective therapy for treatment-resistant depression and post-traumatic stress disorder (Feder et al., 2014; Walsh et al., 2019; Shiroma et al., 2020). While high anesthetic dose of ketamine can be neurotoxic (Morris et al., 2021), recent studies have shown that ketamine is neuroprotective in rodent models of cerebral ischemia and traumatic brain injury and having anti-inflammatory effects via inhibition of both microglia and astrocyte activation (Shu et al., 2012; Wang et al., 2017; Chen et al., 2017). Additionally, deep brain stimulation has been shown to increase BDNF in the nigrostriatal system and also has a neuroprotective effect for dopaminergic neurons in the SNc (Maesawa et al., 2004; Spieles-Engemann et al., 2010; Fischer et al., 2017). Our group has previously shown that increased BDNF-signaling via striatal TrkB receptors underlies the long-term anti-dyskinetic activity of ketamine (Bartlett et al., 2020). In the current study, we show that BDNF is not involved in the neuroprotective effects of ketamine treatment. We first conducted behavioral tasks in which we evaluated parkinsonian phenotypes, the amphetamine-induced rotation test as a measure of parkinsonian lesion severity and the forelimb adjusting steps test as a measure of parkinsonian akinesia, and tested ketamine and the TrkB receptor antagonist ANA-12. We then evaluated changes in dopaminergic neuron count, *TrkB* RNA, tyrosine hydroxylase and p-ERK levels in the striatum and substantia nigra to examine if sub-anesthetic ketamine treatment was neuroprotective via neurotrophic effects.

### 4.1 TrkB-antagonism did not block the robust motor improvements induced by low-dose ketamine treatment

As our first main finding, we demonstrated that the 1-week daily sub-anesthetic ketamine treatment led to robust motor improvements 28 days post-lesion in this progressive mild 6-OHDA-lesion model, but that the TrkB receptor antagonist ANA-12 did not block the protective activity of ketamine treatment (**Figure 2**). The amphetamine-induced rotation test is used as a measure of lesion severity with more rotations made ipsilateral to the lesioned hemisphere indicating more dopamine denervation. Ketamine treatment significantly decreased the number of ipsilateral rotations compared to vehicle at day 28 post-treatment, indicating a protective effect. Co-treatment of ketamine + ANA-12 surprisingly did not block the effect of ketamine but rather decreased the number of ipsilateral rotations compared to the vehicle group even further (**Figure 2B**), indicating that the TrkB receptor axis is not involved in the neuroprotective activity of ketamine that led to robust motor improvement. The forelimb adjusting steps (FAS) test is used as a measure of parkinsonian akinesia, with less steps made by the animal indicating more dopamine neuron denervation (Schallert et al., 1979; Olsson et al., 1995). The low-dose ketamine-treatment significantly increased mean number of steps made by the animals, reducing the parkinsonian akinesia phenotype almost completely, as shown in **Figure 2C**. It is important to point out that in Bartlett et al., 2020, we had shown that low-dose ketamine also improves parkinsonian motor symptoms acutely in the established unilateral 6-OHDA-lesioned rat model of PD, both increasing stepping in the FAS test and performance on the RotaRod. Yet here we are testing one and three weeks after the last ketamine exposure, so are using this as non-drug induced behavior test to evaluate lesion severity to complement the amphetamine-induced rotation test. And again, surprisingly the ANA-12 treatment did not block the effect of ketamine, further indicting neuroprotective activity that is independent of neurotrophic activity by BDNF signaling. And since we blocked BDNF signaling during the ketamine treatment this can neither be due to BDNF signaling induced plasticity, in contrast to what we had shown for the long-term anti-dyskinetic activity of low-dose ketamine that was completely blocked by ANA-12 (Bartlett et al., 2020).

### 4.2 Effects of ketamine and ANA-12 in the substantia nigra pars compacta and striatum

The second main finding of this study is that ketamine treatment does not lead to significant protection of dopaminergic neurons in the SNc (**Figure 3A**) compared to vehicle, as there were no differences in remaining dopamine neuron count post-lesion between any of the treatment groups. Interestingly, the K + ANA-12 showed a trend of an increase in the remaining TH-positive neuron number on the lesioned hemisphere (∼30% increase vs. vehicle). We validated successful target engagement of the TrkB receptor with ANA-12, by the confirmation that there was a significant decrease in *TrkB* RNA in both the ipsilateral and contralateral hemispheres of the substantia nigra in the K + ANA-12 group vs. ketamine-only (**Figure 4**). These findings suggest that there is no effect of ketamine in the substantia nigra to promote neuroprotective effects.

The third main finding in the current study is in the evaluation of changes in striatal tyrosine hydroxylase (TH) levels. We have shown that there is a trend of a small increase in striatal TH level in the lesioned striatum in ketamine treated animals compared to the vehicle group. This indicates that ketamine may have some but only marginal protective activity for dopaminergic terminals in the striatum rather than for the cell bodies in the substantia nigra. Again, this increase was surprisingly not blocked but further increased in the K + ANA-12 group, where a significant increase of striatal TH level vs. both the vehicle and the ketamine-only groups was reached (**Figure 5A**). Yet, there were no significant changes in p-ERK1/2 levels between the vehicle and ketamine treated groups in either the intact or lesioned hemisphere, but with the addition of ANA-12, there was a significant reduction of p-ERK1/2 levels relative to total ERK1/2 levels in the striatum (**Figure 5B**). This also indicates that we did have proper target engagement of ANA-12 at the TrkB receptor, as ERK1/2 regulation is downstream of this receptor in the striatum, and further validation that the neuroprotective effects exhibited by ketamine in the lesioned hemisphere may not be mediated by the TrkB receptor.

Future studies investigating the dopamine transporter (DAT; a transporter protein integral in synaptic clearance of dopamine & surrogate index of increased dopamine release) and its level in the striatum will be of importance to begin to understand the more complex mechanisms related to ketamine’s protective motor effects, and to determine if there was evidence of upregulated dopamine-activity in remaining nigrostriatal terminals following ketamine administration, suggesting an enhanced dopamine-tone which is not mediated by TrkB signaling. In reward areas esketamine or ketamine treatment was shown prior to indeed lead to increases in the dopamine-tone in the nucleus accumbens (Rizzo et al., 2025), or the medial prefrontal cortex (Bonaventura et al., 2021).

There are limitations to the current study reported here. We did this work to start assessing the mechanism of neuroprotective effects of ketamine only in male rats, since the anti-parkinsonian effects of ketamine had not yet been investigated in females, and future studies should investigate if there is any sex-specificity. We also focused on the hypothesis that this effect might be driven by neurotrophic BDNF signaling. Yet, in addition to the BDNF-driven effects, there is also evidence for ketamine having anti-inflammatory effects via inhibition of both microglia and astrocyte activation (Shu et al., 2012; Wang et al., 2017; Chen et al., 2017). Studies show that microglial activation is a major contributor to degeneration seen in PD (Block et al., 2007; Kannarkat et al., 2013) and that astroglia might be involved in neuroprotective activity of ketamine in a PD model via inhibition of the AKT kinase axis and promotion of autophagy (Xiong et al., 2024). In a 6-OHDA rodent model of LID, ketamine was shown to increase striatal interleukin-6 (IL-6) in dyskinetic rats, indicating a modulation of inflammatory response (Pottenger et al., 2021). A similar increase in IL-6 after ketamine was also seen in plasma of patients in an open label dose finding trial in individuals with LID (Falk et al., 2023). Additionally, it was also shown that there was an increase in CD68 in the lesioned SNc hemisphere after ketamine treatment, indicating increased microglia activation through the phagocytic pathway (Pottenger et al., 2021). Ketamine has also been shown to inhibit the production and release of other pro-inflammatory cytokines such as IL-1beta and TNF-alpha in macrophages and both microglia and astrocytes in rodents (Chang et al., 2009; Chang et al, 2010; Yuhas et al., 2015). The protective effects of ketamine could, therefore, instead also be in part mediated by anti-inflammatory mechanisms.

### 4.5 Conclusion

In conclusion, we show that ketamine at the low dose currently pursued for long-term therapeutic benefit for individuals with LID (Falk & Cowen, 2026) is not neurotoxic but rather shows some protective activity in a standard preclinical model of PD degeneration. The protective activity is not driven by neurotrophic activity, rather indicating increased dopamine tone or immunomodulatory actions as possible contributors. This is an important safety aspect for the ongoing clinical development for sub-anesthetic ketamine treatment for individuals with LID, in light of a prior retrospective case study showing anti-dyskinetic activity of low-dose ketamine infusion (Sherman et al., 2016), and the recently reported encouraging safety and tolerability data from an open label dose-finding Phase I clinical trial (Falk et al., 2023).

## Acknowledgements

We would like to thank the following funding sources: NIH-NINDS R56 NS109608 & R01 NS122805 (TF); Arizona Biomedical Research Commission (ABRC) ADHS18-198846 (SJS, TF); and the Mary Mack Estate gift.

## Declaration of Competing Interest

SJS and TF have a patent for the use of ketamine as a novel treatment for levodopa-induced dyskinesia associated with Parkinson’s disease that was licensed to PharmaTher Inc. 2020-2026. They have consulted for PharmaTher Inc. in 2021 and 2023. TF also received travel support. Mitchell J. Bartlett owns stock in PharmaTher Inc. These interests did not influence the study design, data collection, analysis, interpretation, manuscript preparation, or decision to publish. All other authors declare no competing financial or personal interests.

## CRediT authorship contribution statement

**Carolyn J. Stopera**: Formal analysis, Investigation, Methodology, Validation, Writing – original draft. **Mitchell J. Bartlett**: Formal analysis, Investigation, Methodology, Supervision, Validation. **Kelsey Bernard**: Investigation, Methodology. **Rachel Conklin**: Investigation, Methodology. **Sansita Singh**: Investigation, Methodology. **Jennifer B. Frye**: Investigation, Methodology. **Lalitha Madhavan**: Resources, Methodology. **Kristian R. Doyle**: Resources, Methodology. **Scott J. Sherman**: Writing – review & editing. **Kathy Steece-Collier**: Writing – review & editing. **Torsten Falk**: Conceptualization, Funding acquisition, Resources, Supervision, Writing – review & editing.

